# Regulation of polyphosphate homeostasis in *S. cerevisiae* by the Sky1 protein kinase

**DOI:** 10.64898/2026.09.24.753767

**Authors:** Alix Denoncourt, Sarah McKeague, Selin Gabbarizadeh, Michael Downey

## Abstract

Eukaryotic cells store inorganic phosphate in the form of long chains called polyphosphates (polyP) that can reach many hundreds of residues in length. In the budding yeast *Saccharomyces cerevisiae*, polyP is synthesized by the VTC complex using ATP as a substrate and sequestered at high concentrations in the vacuole. Here, polyP chain length and turnover are regulated by the polyphosphatase enzymes Ppn1 and Ppn2. In the vacuole lumen, polyP plays a critical role in phosphate and ion homeostasis. Beyond the vacuole, minor populations of polyP are thought to impact various cellular functions, including in the nucleus, in part by binding to target proteins. We speculated that polyP metabolism must be tightly coupled to mechanisms of homeostatic control operating elsewhere in the cell. In this work, we report that cells lacking Sky1, a serine-arginine repeat protein kinase, have very low levels of polyP, an effect that is exacerbated by alkaline stress. Under these conditions *sky1*Δ mutants also display increased expression of a subset of genes from the PHO regulon. Genetic analysis suggests that both phenotypes are linked to an increase in potassium influx via the Trk1 potassium transporter. Loss of Npl3, a known target of Sky1, partially rescues the phosphate homeostasis defects in *sky1*Δ mutants, although this occurs independently of a well characterized Sky1 phosphorylation site on Npl3. Finally, the loss of polyP in *sky1*Δ mutants depends on the Ppn1 and Ppn2 polyphosphatases and the Pep4 protease located in the vacuole. Altogether, our work provides novel insights into how polyP homeostasis in the vacuole is coordinated with events occurring throughout the cell.

## IMPORTANCE

In diverse types of cells, inorganic phosphate can be assembled into long chains called polyphosphates. In addition to acting as a phosphate reserve, these polyP chains, which can reach hundreds of residues in length, engage in important functions throughout the cell. How cells coordinate polyP synthesis and turnover in step with events occurring elsewhere in the cell remains poorly understood. Under standard laboratory conditions, cells of the budding yeast *S. cerevisiae* accumulate high levels of polyP in their vacuoles. For this reason, yeast has emerged as a premiere model for understanding fundamental aspects of polyP biology. In this work, we leverage a screen for new genes that impinge on polyP metabolism to identify the conserved Sky1 protein kinase as required for normal polyP accumulation. We characterize Sky1 as a central player in a complex network at the centre of ion homeostasis at the plasma membrane, gene expression in the nucleus, and polyP turnover in the vacuole. Altogether, our work serves as a foundation on which we can better understand how polyP metabolism is coordinated in space and time within the cell.

## INTRODUCTION

Polyphosphates (polyP) are conceptually simple molecules consisting of long chains of inorganic phosphates joined by high energy bonds^1^. They are found in diverse prokaryotic, eukaryotic, and archaeal cells, although the mechanism of synthesis is species specific. In the budding yeast *S. cerevisiae*, polyP synthesis depends on the action of the VTC complex consisting of Vtc4 (the catalytic subunit), Vtc1, and either Vtc2 or Vtc3^2^. VTC spans the vacuolar membrane and coordinates the transfer of the gamma phosphate of cytoplasmic ATP to a growing polyP chain, coincident with the translocation of that chain into the vacuole lumen^3^. Here, polyP accumulates to high concentrations that can reach over 200 mM (measured as phosphate monomers)^4^.

In yeast, a recently updated model posits that levels of a particular inositol pyrophosphate species, called IP_8_, drop during phosphate starvation^5^. This drop in IP_8_ levels promotes Pho81 repression of the Pho85-Pho80 CDK-cyclin pair^5^, resulting in downstream translocation of the Pho4 transcription factor to the nucleus to promote the expression of >20 genes^5–7^. These genes comprise the ‘PHO’ regulon and include those associated with phosphate uptake into the cell (*e.g.* genes encoding the membrane transporters Pho84 and Pho89) and phosphate scavenging (*e.g.* the gene encoding the Pho5 phosphatase that acts on phosphate monoesters at the cell surface)^6^. Subunits of the VTC complex, whose activity is promoted by IP_8_, are also upregulated during phosphate starvation^6^, presumably to prime the cell for rapid polyP synthesis when inorganic phosphate (Pi) becomes available. An important outcome of polyP sequestration in the vacuole is that cytoplasmic Pi levels are kept low, which promotes continued uptake from the environment. Inside the vacuole lumen, the dual exo/endopolyphosphatase Ppn1 and endopolyphosphatase Ppn2 process polyP to generate chains of mixed length^8, 9^. These enzymes are also involved in mobilizing polyP stores to provide free inorganic phosphate during starvation. Little is known about the molecular control over Ppn1 and Ppn2 activities, although they are both expressed as part of the PHO regulon^7^. Recent work suggests that Ppn1 and Ppn2 are negatively regulated by free phosphate in the vacuole^10^. Additionally, Ppn1 activation requires proteolytic cleavage by Pep4 and other vacuolar proteases^11^. Two additional polyphosphatases, Ppx1 and Ddp1, are localized to the cytoplasm and cytoplasm/nucleus respectively^12^, although their direct role in polyP metabolism, if any, is unclear.

In mammalian cells, measured levels of polyP are highly variable, with reported concentrations dependent on the techniques used for extraction and analysis^13^. Surprisingly, the mechanism of polyP synthesis itself is largely unclear^14^. The mitochondrial FoFI ATP synthetase has been proposed to act as a polyP biosynthetic enzyme^15^, but its contribution to total polyP pools in the cell has not been determined. Several polyphosphatase enzymes have been suggested to act in human cells^12^, with the NUDT3 endopolyphosphatase (a homolog of Ddp1^16^) thought to play a role in regulating polyP accumulation in the nucleus during oxidative stress^17^.

It is becoming increasingly clear that polyP has broad functions throughout the cell. For example, yeast *vtc4*Δ mutants have defects in polysome assembly, indicative of a role in translation control or ribosome biogenesis^18^. PolyP is also proposed to function in ion homeostasis^19^, cell cycle control and genome stability^20^. Roles for polyP in processes occurring beyond the vacuole could be indirect consequences of its role in phosphate homeostasis, but more direct roles are also possible. For instance, it is noteworthy that minor populations of polyP are found outside the vacuole^21^, including in the nucleus^22^, and polyP can bind to proteins involved in transcriptional regulation and ribosome assembly^18^.

It stands to reason that polyP metabolism should be coordinated with events across the cell, but the details of this molecular crosstalk remain poorly understood. Here, we illuminate a hitherto unknown connection between the Sky1 protein kinase, ion transport at the plasma membrane, and the regulation of polyP hydrolysis during alkaline stress. Altogether, this work provides novel insights into how polyP metabolism is linked to cell signalling pathways operating beyond the vacuole.

## RESULTS

We were motivated to identify novel pathways that regulate polyP accumulation and surmised that polyphosphate homeostasis may be regulated by enzymes that catalyze post-translational modifications such as phosphorylation. Therefore, we screened 112 non-essential kinase mutants from the yeast haploid deletion set^23^ for changes in polyP levels (**Fig. 1A**). Many gene deletions showed either small changes in polyP or changes that were not consistent across multiple independent experiments. Two gene mutants stood out in multiple trials as having profound effects on polyP accumulation: *pho85*Δ (increased polyP) and *sky1*Δ (decreased polyP) (**Fig. 1B**). Note that in these gels RNA species extracted with polyP form distinct bands at the top of the gel and serve as a loading control (**Fig. 1B**). Pho85 encodes a cyclin-dependent protein kinase that works with the Pho80 cyclin to repress the PHO regulon during phosphate replete conditions. Deletion of either *PHO85* or *PHO80* results in hyperactivation of the PHO regulon during standard growth conditions^24, 25^. The increase in polyP in *pho85*Δ mutants was described previously by several groups^26, 27^. Curiously, earlier work by Ogawa *et al.* described a *pho80*Δ strain as having less polyP than wild-type cells^7^, even though these deletions would be expected to impact polyP in the same way. While our analysis of the *pho80*Δ strain from the deletion collection indicated increased polyP relative to the wild-type control, the amount of polyP was still less than that observed in *pho85*Δ mutants (**Fig. S1A**). Re-generation of *pho80*Δ mutants in the same genetic background in our own lab showed an accumulation of polyP in line with what we observed for *pho85*Δ (**Fig. S1B**). Based on this observation, we surmise that increased expression of PHO-regulated genes such as *PHO84* and *VTC* are together responsible for the hyperaccumulation of polyP in *pho85*Δ and *pho80*Δ cells. The *pho80*Δ mutant from the deletion set likely contains one or more extragenic mutations that partially reverse the polyP hyperaccumulation phenotype. To rule out confounding effects of suppressor mutations, yeast strains described in the rest of the study were made *de novo*, and key phenotypes were confirmed by rescuing mutations with plasmid-borne copies of deleted genes.

**Figure 1.**
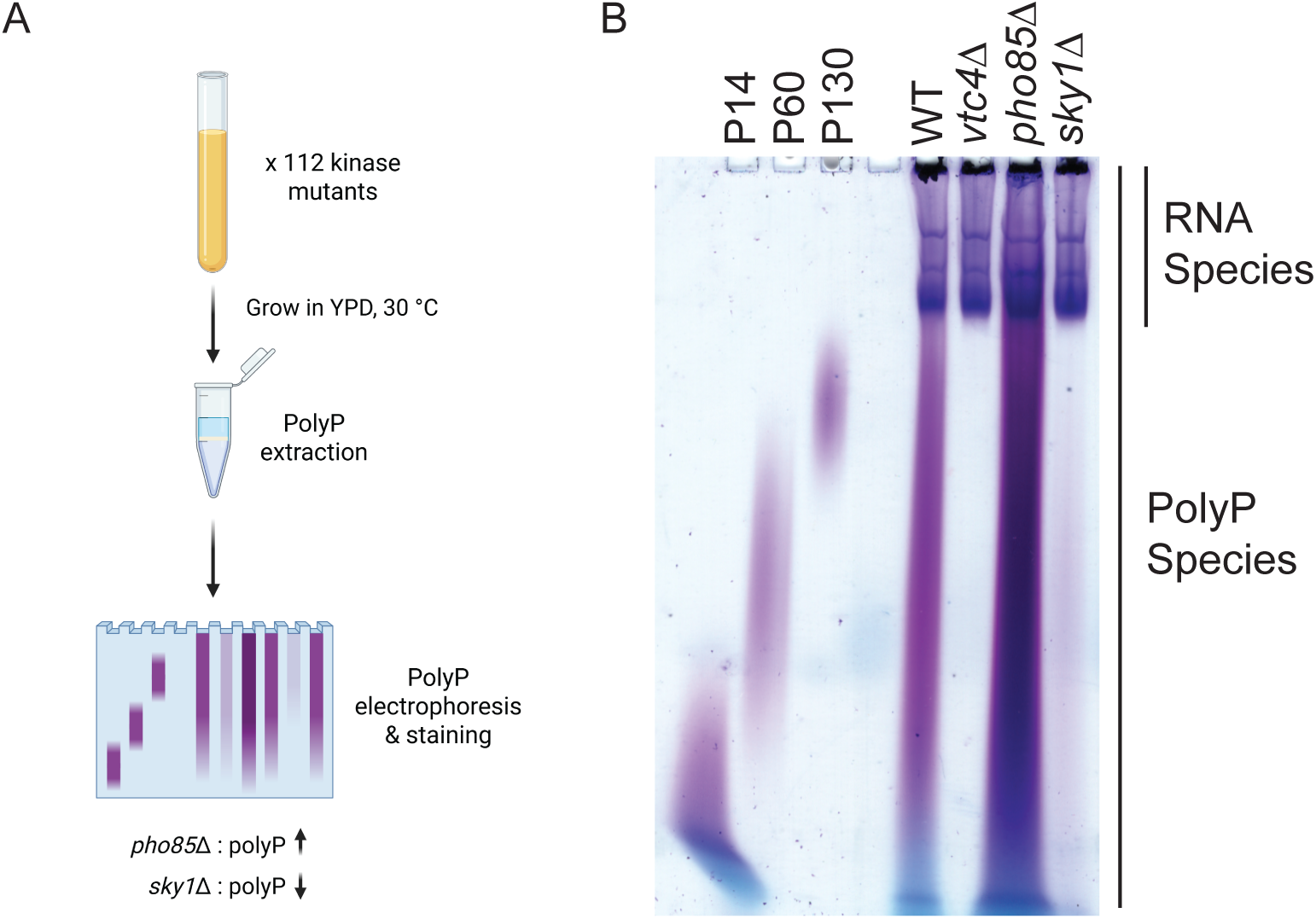
Screen of the yeast haploid deletion set uncovers altered polyP levels in *pho85*Δ and *sky1*Δ mutants. **(A)** A schematic representation of the polyP extraction protocol used in this work. Image was made using Biorender. **(B)** PolyP was extracted from cells grown to exponential phase in YPD media and stained with toluidine blue following TBE-urea PAGE separation. Image is representative of data from ≥ 3 experiments.

In contrast to *pho85*Δ, *sky1*Δ mutants had very low levels of polyP (**Fig. 1B**). Quantification of these levels using the *in vitro* ‘Phosfinity Quant’ assay^28^ showed 10-fold less polyP compared to wild-type controls (**Fig. 2A**). Unexpectedly, the loss of polyP in *sky1*Δ mutants relative to wild-type controls was more apparent when cells were grown in YPD media versus synthetic complete (SC) media (**Fig. 2B**). Analysis of the literature suggested a connection between Sky1 and intracellular pH. Specifically, although the relevant direct target is unknown, Sky1 has been proposed to negatively regulate potassium import across the plasma membrane by inhibiting the Trk1/Trk2 potassium transporters^29–31^. Therefore, potassium import increases in *sky1*Δ mutants because Trk1/Trk2 are hyperactive^31^. As intracellular potassium rises, the Pma1 ATPase is proposed to export H+ ions to compensate for increased positive charge on the inside of the plasma membrane, resulting in an elevated intracellular pH and membrane depolarization^32–35.^

**Figure 2.**
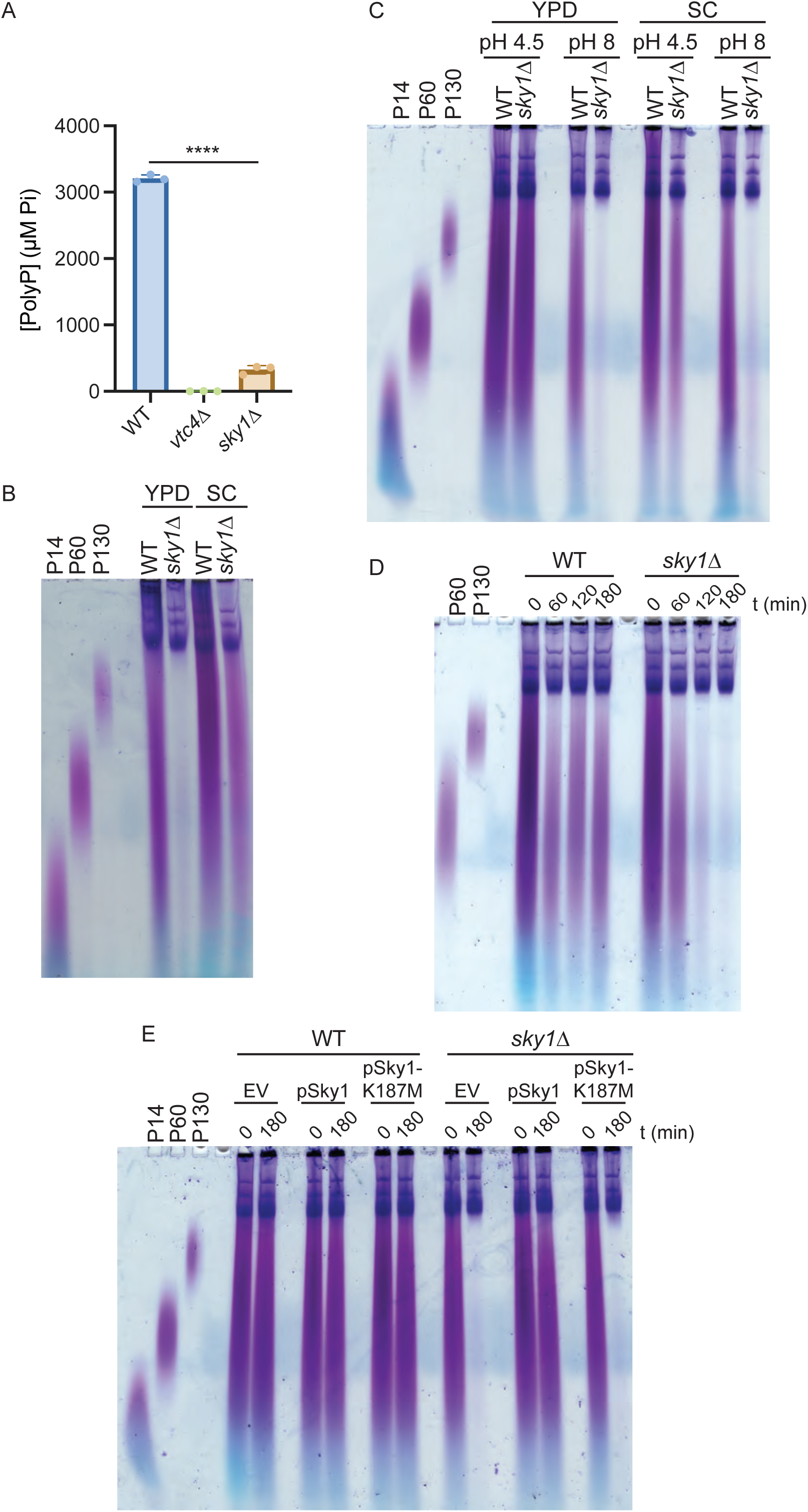
PolyP levels in *sky1*Δ are modulated by variations in pH. **(A)** PolyP was extracted from the indicated strains grown to exponential phase in YPD media and quantified using the *in vitro* Phosfinity polyP quantification assay. Data are presented as means ± SD. Significance was determined by ordinary one-way ANOVA with Tukey’s multiple comparisons test. ****, *p* < 0.0001. Significance is indicated only for key comparisons discussed in the text, with all *p* values presented in **Supplementary Table S5**. **(B)** WT and *sky1*Δ cells were grown to exponential phase in YPD or synthetic complete (SC) minimal media, and polyP was extracted and visualized via toluidine blue staining following TBE-urea PAGE separation. **(C)** WT and *sky1*Δ cells were grown to exponential phase in YPD or SC media with pH adjusted to 4.5 or 8. PolyP was extracted and visualized on gel as described above. **(D)** Cells were grown in YPD pH 4.5 for approximately 1 cell division then switched to YPD pH 8. Cells were collected before the switch and at the indicated time points after the switch to alkaline conditions for polyP extraction and visualization as described above. **(E)** WT and *sky1*Δ cells bearing an empty vector or a vector expressing the wild-type or K187M *sky1* allele were grown in YPD pH 4.5 for approximately 1 cell division then switched to YPD pH 8. Cells were collected before the switch and after 3 hours in alkaline conditions for polyP extraction and visualization as described above. Cells in **(E)** are all in the *PHO5*-3HA background (see text describing Figure 4). Images are representative of data from ≥ 3 experiments.

Previous work demonstrated rapid polyP loss in response to extracellular alkaline stress, which was proposed to be important for pH buffering in the vacuole^36, 37^. Therefore, we reasoned that differences in polyP levels across media types could stem from differences in pH. Our YPD has a pH of 6, whereas our synthetic complete media has a pH of 4.5. We tested the effect of high and low pH on polyP accumulation in both media types. Cells grown at pH 4.5 had higher levels of polyP than at pH 8, regardless of the media type used (**Fig. 2C**), making the impact of *sky1*Δ easier to discern. To further characterize this phenomenon, we carried out media switch experiments wherein strains were first grown in YPD at pH 4.5 before a shift to YPD at pH 8 for 3 hours. In this experiment, wild-type polyP decreased before stabilizing at the end of the time course, consistent with what was described previously^37^ (**Fig. 2D**). The *sky1*Δ mutant cells underwent the same decrease in polyP but failed to stabilize in the same manner as their wild-type counterpart (**Fig. 2D**). To test if the Sky1 kinase activity was required for its function in polyP homeostasis, we used these media switch experiments to analyze a *sky1*Δ strain transformed with a vector expressing the wild-type *SKY1* allele, a vector expressing a previously described kinase-dead mutant allele (K187M)^38^, or an empty vector control. The *sky1*Δ polyP phenotype was fully rescued by plasmid-based expression of *SKY1* under its native promoter, but not by the kinase-dead K187M allele (**Fig. 2E**), confirming dependence on kinase activity.

We next investigated the genetic dependencies of the *sky1*Δ polyP phenotype. Since increased potassium import in *sky1*Δ mutants depends on Trk1 and Trk2^31^ (**Fig. 3A**), we might expect to reverse the polyP accumulation defects in *sky1*Δ mutants by deleting the genes encoding these proteins. While we were unable construct *sky1*Δ *trk1*Δ *trk2*Δ triple mutants, *TRK1* deletion reversed the polyP loss in *sky1*Δ mutants under alkaline conditions (**Fig. 3B**). Our ability to observe an impact of *trk1*Δ without concomitant deletion of *TRK2* is consistent with previous work suggesting a more prominent role for Trk1 in potassium transport compared to Trk2^39^. Notably, potassium import by Trk1 (and Trk2) is thought to be negatively regulated by the Ppz1 and Ppz2 phosphatases^29, 40^ (**Fig. 3A**). Deletion of both *PPZ1* and *PPZ2* also decreased polyP levels relative to wild-type controls (**Fig. 3B**), consistent with previous reports that these mutants have increased potassium levels and a depolarized plasma membrane^34^.

**Figure 3.**
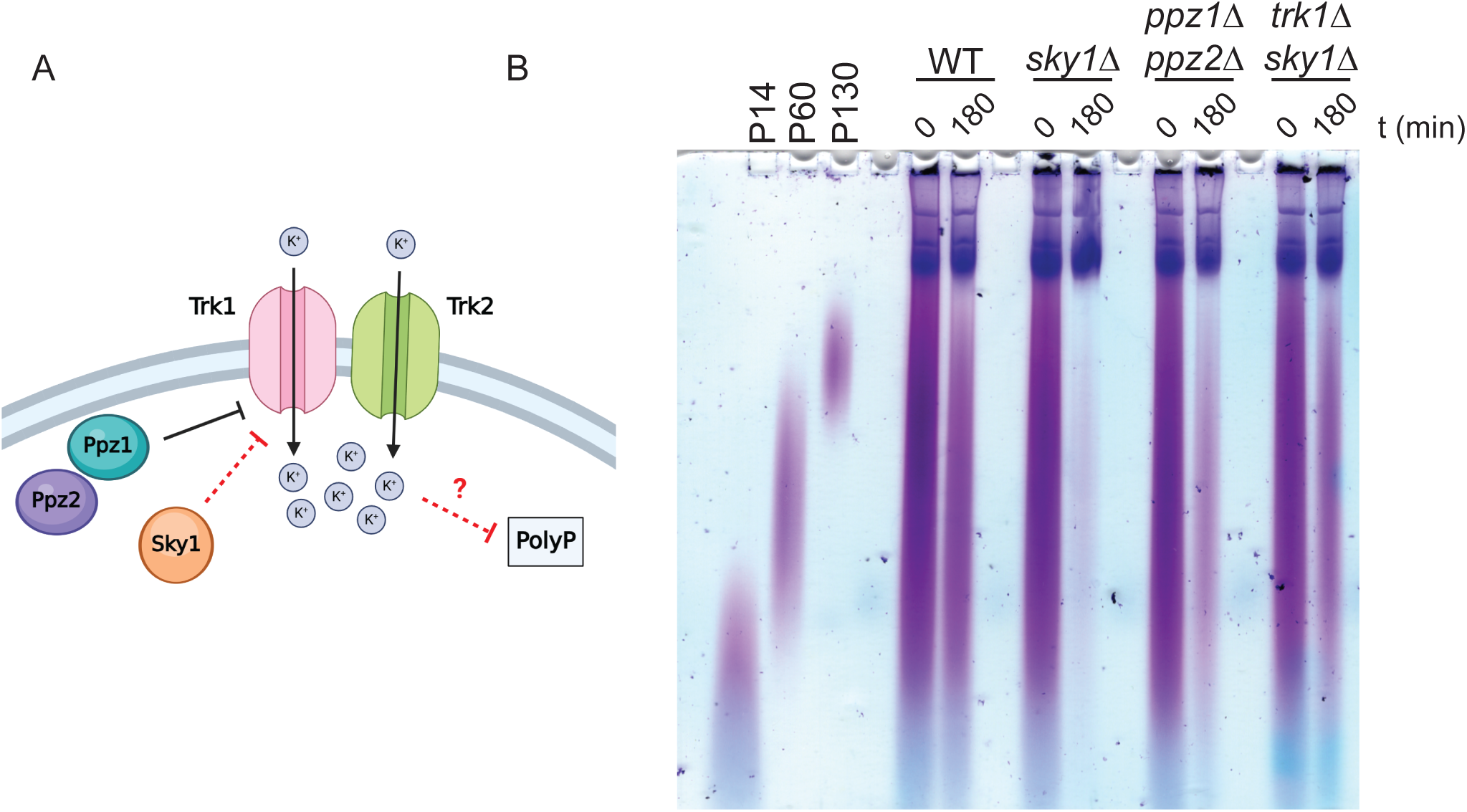
Gene inactivation of the Trk1 potassium importer rescues polyP levels in *sky1*Δ during alkaline stress. **(A)** Schematic model illustrating regulation of the Trk1/2 transporters by the Ppz1 and Ppz2 phosphatases. Sky1 is thought to negatively regulate potassium uptake directly or indirectly via Trk1/2. Image was made using Biorender. **(B)** Indicated strains were grown in YPD pH 4.5 for approximately 1 cell division then switched to YPD pH 8. Cells were collected before the switch and after 3 hours in alkaline conditions for polyP extraction, TBE-urea PAGE separation, and toluidine blue staining. Image is representative of data from ≥ 3 experiments.

Alkaline stress was also shown to activate the phosphate starvation response in a Pho4-dependent manner^37, 41^, although the details of this regulation are poorly understood. To investigate PHO regulation in *sky1*Δ mutants, we examined the expression of the acid phosphatase Pho5, a marker of PHO activation. In these experiments *PHO5*-3HA is expressed from the native chromosomal location. Expression of the reporter was induced by alkaline stress in wild-type cells, and this induction was much more dramatic in *sky1*Δ mutants (**Fig. 4A**). Elevated expression of Pho5-3HA in *sky1*Δ mutants was fully dependent on the canonical PHO pathway since deleting *PHO4* led to a complete Pho5-3HA repression (**Fig. 4A**). Like polyP levels, this molecular phenotype was rescued by plasmid-based expression of *SKY1* but not the K187M allele encoding the kinase-dead protein (**Fig. 4B**). Notably, wild-type cells transformed with pAG36-*SKY1* strongly repressed Pho5-3HA compared to those bearing the empty vector, consistent with Sky1 acting as an overall negative regulator of Pho5-3HA expression (**Fig. 4B**). Like the polyP phenotype, the increased expression of Pho5-3HA in *sky1*Δ was rescued by deletion of *TRK1* (**Fig. 4C**). How PHO induction in *sky1*Δ mutants relates to the decrease in polyP is not clear. Indeed, we expected that derepression of the PHO regulon would lead to an increase in polyP accumulation, as seen in *pho85*Δ and *pho80*Δ mutants (**Fig. S1**). Moreover, despite having a *sky1*Δ-like phenotype in terms of polyP accumulation (albeit not as dramatic), the *ppz1*Δ *ppz2*Δ mutant showed decreased Pho5-3HA expression relative to wild-type controls (**Fig. 4C**).

**Figure 4.**
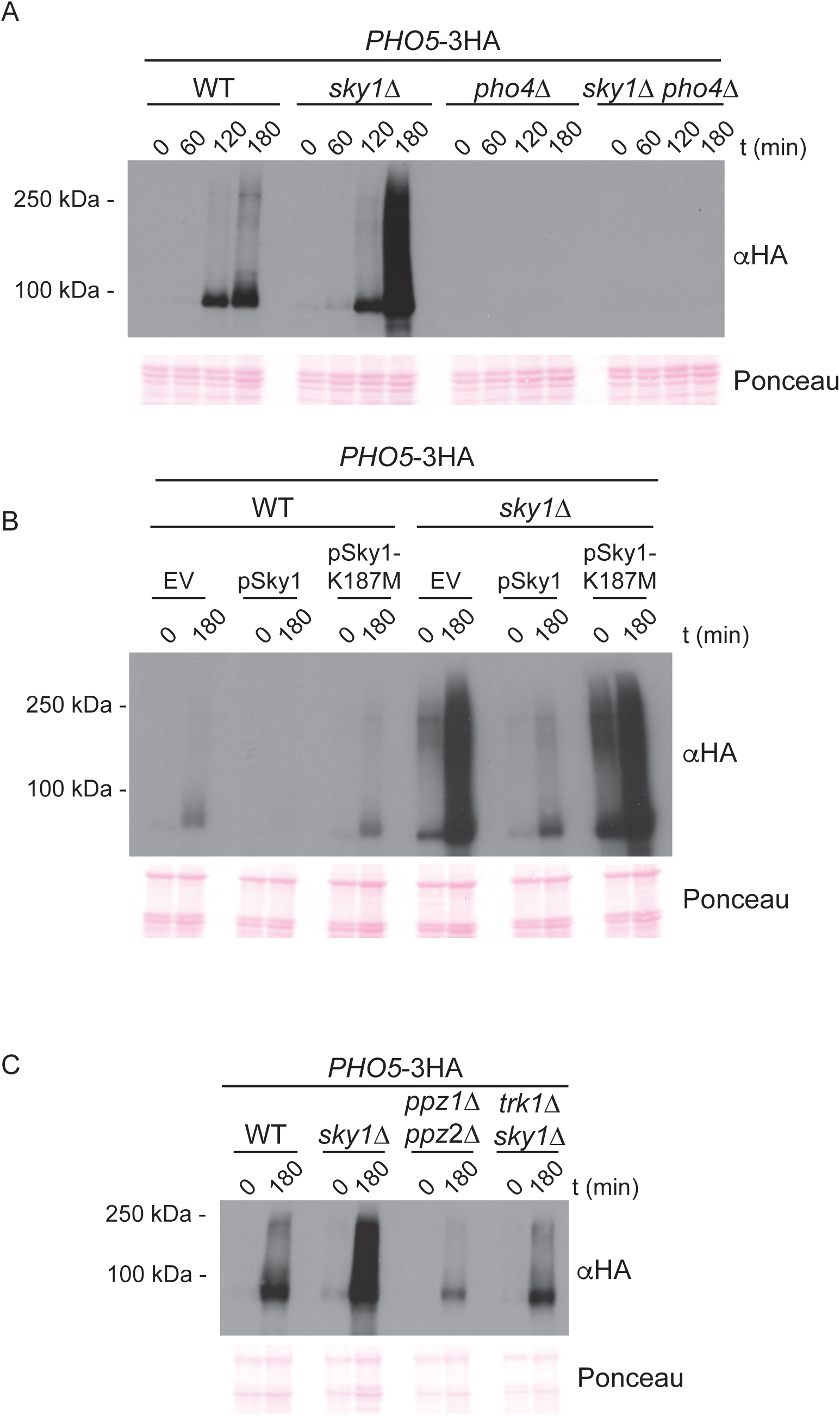
*sky1*Δ mutants show increased accumulation of the acid phosphatase Pho5 compared to wild-type cells during alkaline stress. **(A)** to **(C)** Indicated strains expressing *PHO5-*3HA from the chromosomal locus were grown in YPD pH 4.5 for approximately 1 cell division then switched to YPD pH 8. Cells were collected at the indicated time points for protein extraction, SDS-PAGE separation, and western blotting. Pho5-3HA protein was detected using an anti-HA antibody. Ponceau S staining of the membrane is shown as loading control. Images are representative of data from ≥ 3 experiments.

Sky1 was first characterized by the Guthrie Lab as a serine-arginine (SR)-repeat protein kinase that regulates the nuclear accumulation of Npl3, involved in mRNA export and splicing^38, 42^. Subsequent studies identified the RNA-binding proteins Gbp2 and Hrb1 as additional targets of Sky1, although the *in vivo* relevance of Hrb1 phosphorylation is unclear^43, 44^. In an effort to identify the missing link between Sky1, potassium homeostasis, and polyP regulation, we next assessed polyP accumulation in mutants deleted for known targets of Sky1. While *gbp2*Δ and *hrb1*Δ single mutants showed no obvious change in polyP levels (**Fig. S2B**), deletion of *NPL3* resulted in increased polyP levels which was particularly evident after 1 hour in alkaline conditions (**Fig. 5A & Fig. 5B**). Deleting *NPL3* in the *sky1*Δ background gave a polyP phenotype profile intermediate between wild-type and *sky1Δ,* and allowed for retention of some polyP after 180 minutes (**Fig. 5A & Fig. 5B**). We also generated a quadruple mutant where genes encoding all three Sky1 targets – *NPL3*, *GBP2* and *HRB1* – were deleted in the *sky1*Δ background. This quadruple mutant behaved like *sky1*Δ *npl3*Δ (**Fig. S3A**), indicating that Npl3 is the main driver of this polyP phenotype. Notably, the loss of *NPL3* also reversed hyperaccumulation of *PHO5* mRNA and Pho5-3HA protein in *sky1*Δ mutants during alkaline stress (**Fig. 5C & Fig. S3B**). Other PHO genes showed a varied response. *PHO84* mRNA followed a similar trend to that observed for *PHO5* (**Fig. S3B**). In contrast, *PHO89* mRNA was increased in *sky1*Δ mutants compared to wild-type controls, but this increase was not impacted by co-deletion of *NPL3* (**Fig. S3B**). These data highlight a complex role for *NPL3* in phosphate homeostasis. Importantly, *npl3*Δ phenotypes could be complemented by *NPL3* expression from a plasmid (**Fig. S4A & Fig. S4B**).

**Figure 5.**
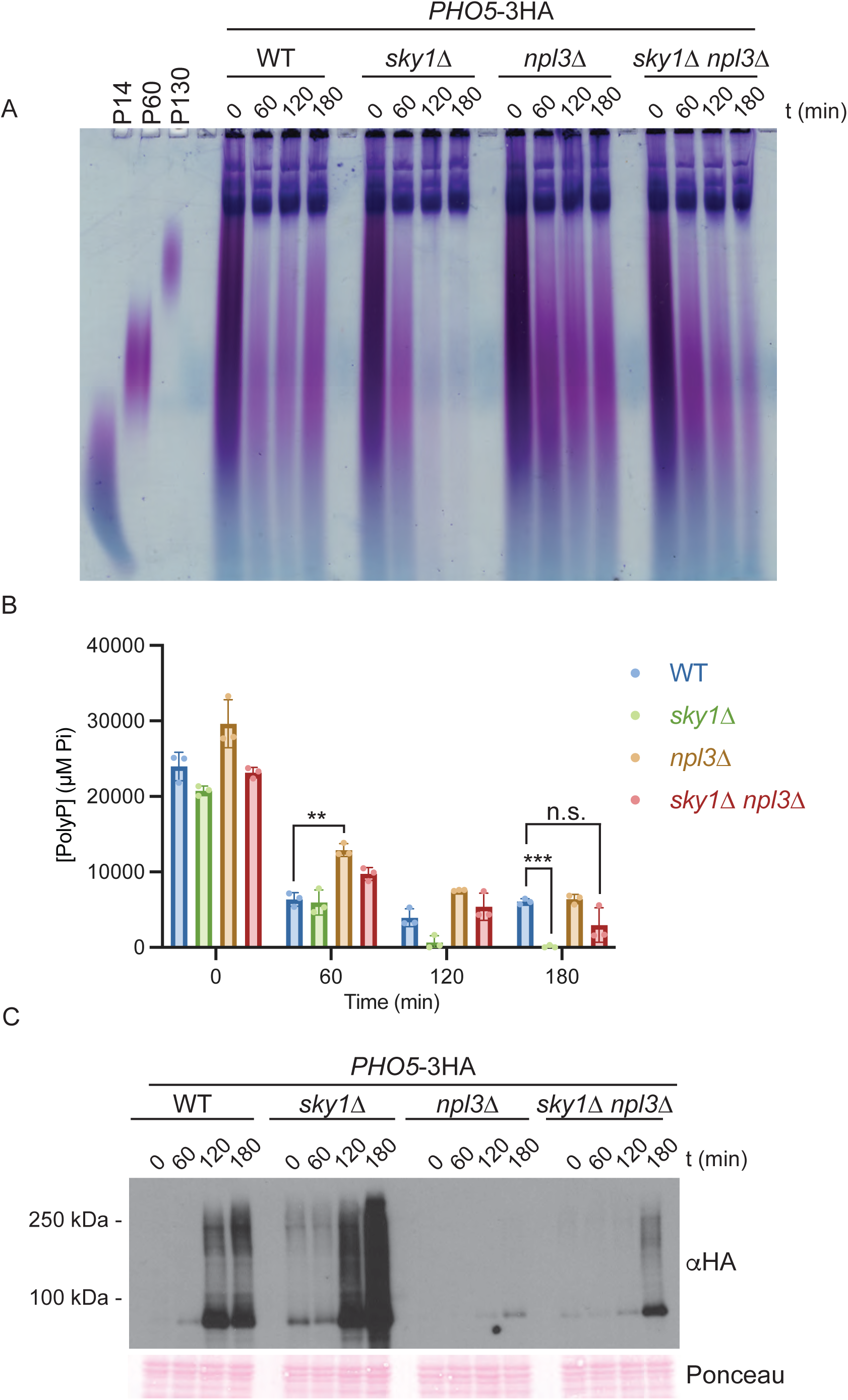
Loss of *NPL3* increases polyP levels and decreases Pho5-3HA protein expression during alkaline stress. **(A)** Indicated strains were grown in YPD pH 4.5 for approximately 1 cell division then switched to YPD pH 8. Cells were collected at 1 hour intervals and polyP was extracted for TBE-urea PAGE separation and toluidine blue staining. **(B)** PolyP samples from **(A)** were quantified using the *in vitro* Phosfinity polyP quantification assay. Data are presented as means ± SD. Significance was determined by ordinary two-way ANOVA with Tukey’s multiple comparisons test. **, *p* < 0.01. ***, *p* < 0.001. n.s., non-significant. Significance is indicated only for key comparisons discussed in the text, with all *p* values presented in **Supplementary Table S5**. **(C)** Cells expressing *PHO5*-3HA from the chromosomal locus were grown as in **(A)** and collected at the indicated time points for protein extraction, SDS-PAGE separation and western blotting. Pho5-3HA protein was detected using an anti-HA antibody. Ponceau S staining of the membrane is shown as loading control. Images are representative of data from ≥ 3 experiments.

While it is tempting to suggest that Sky1 functions through Npl3 to regulate polyP synthesis, the polyP profiles presented in **Fig. 5B**, although complex, are more suggestive of an additive effect rather than the epistatic relationship that we expect from a target-substrate pair. We also note that expression of a mutant form of Npl3 that cannot be phosphorylated on amino acid residue S411, the site targeted by Sky1^42^, does not show polyP or Pho5-3HA expression phenotypes (**Fig. S4A & Fig. S4B**). As discussed below, we favour a model wherein Sky1 impacts polyP homeostasis through diverse and overlapping pathways rather than a single target.

We next tested if polyP loss in *sky1*Δ mutants was driven by the vacuolar endopolyphosphatases Ppn1 and Ppn2. Notably, Ppn2 was previously noted as playing a role in polyP degradation during alkaline stress^45^. As expected^8^, co-deletion of both *PPN1* and *PPN2* resulted in the formation of long polyP chains whose overall levels were difficult to judge based on PAGE analysis alone (**Fig. 6A**). Quantification revealed that polyP levels in non-stressed conditions (YPD at pH 4.5) were slightly lower in *ppn1*Δ *ppn2*Δ double mutants compared to wild-type cells (**Fig. 6B**). While we do not fully understand the reason for this relative decrease, it is possible that long polyP chains serve as a signal to inhibit polyP accumulation under these conditions. Co-deletion of *PPN1* and *PPN2* countered the complete loss of polyP in a *sky1*Δ background (**Fig. 6A & Fig. 6B**). *PPN1* or *PPN2* single deletions in *sky1*Δ did not have the same impact (**Fig. S5A**), indicating that both polyphosphatase enzymes are important for this regulation.

**Figure 6.**
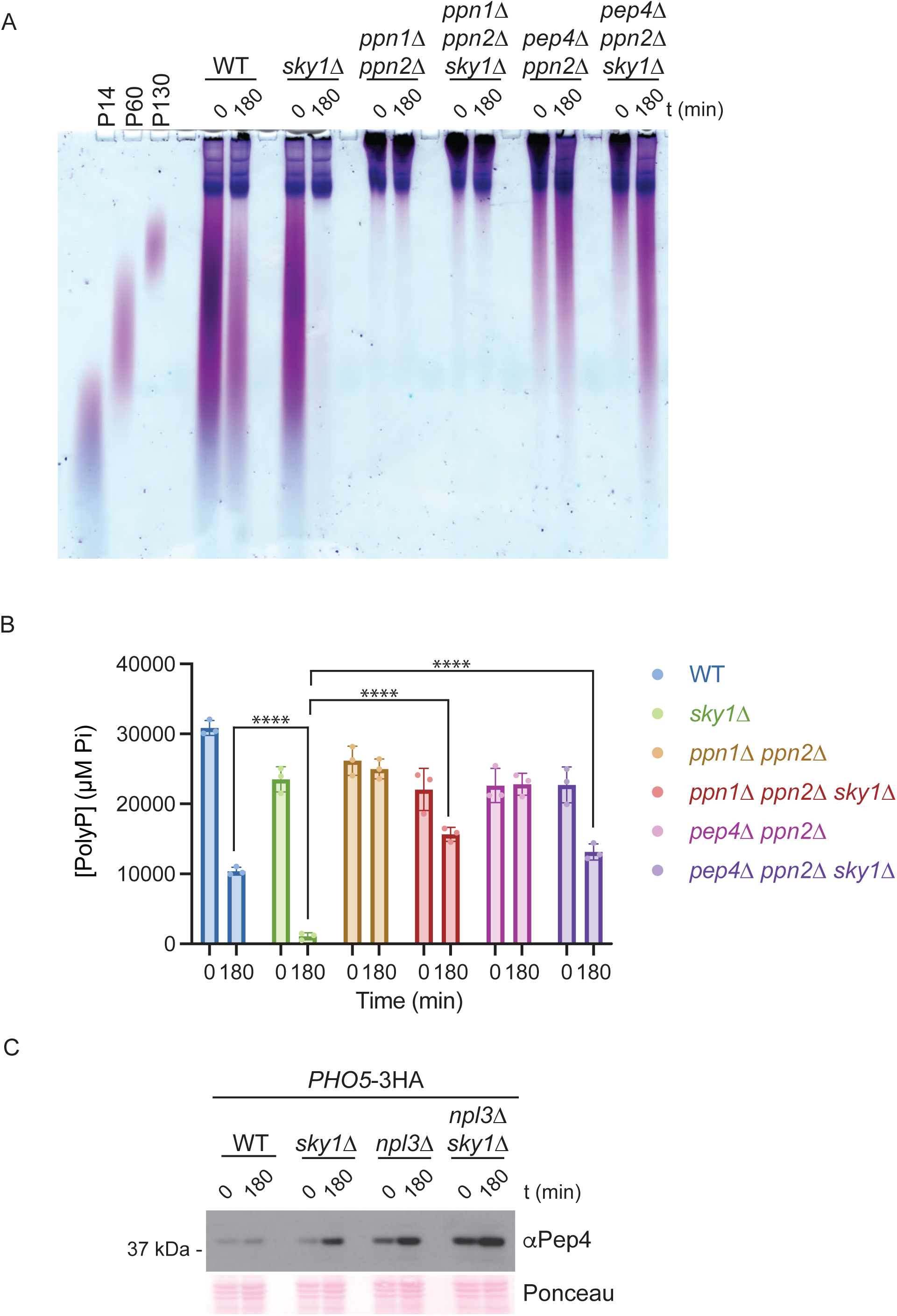
Deletion of the *PPN1* and *PPN2* polyphosphatases in *sky1*Δ mutants prevents loss of polyP in response to alkaline stress. **(A)** Indicated strains were grown in YPD pH 4.5 for approximately 1 cell division then switched to YPD pH 8. Cells were collected before the switch and after 3 hours in alkaline conditions, and polyP was extracted for TBE-urea PAGE separation and toluidine blue staining. **(B)** PolyP samples from **(A)** were quantified using the *in vitro* Phosfinity polyP quantification assay. Data are presented as means ± SD. Significance was determined by ordinary two-way ANOVA with Tukey’s multiple comparisons test. ****, *p* < 0.0001. Significance is indicated only for key comparisons discussed in the text, with all *p* values presented in **Supplementary Table S5**. **(C)** Indicated strains were grown as in **(A)** and cells were collected at 1 hour intervals for protein extraction, SDS-PAGE separation, and western blotting. Membrane was probed with an anti-pep4 antibody. Ponceau S staining of the membrane is shown as loading control. Images are representative of data from ≥ 3 experiments.

This result sparked our interest based on the unanticipated discovery during the course of our work that expression of the Pep4 protease was increased in *sky1*Δ mutants experiencing alkaline stress (**Fig. 6C**). *PEP4* deletion alone did not completely prevent polyP loss in *sky1*Δ cells (**Fig. S5B**), nor did it reverse the Pho5-3HA expression phenotype (**Fig. S5C**). However, Pep4 is a master regulator of protease activities which are together proposed to be required for Ppn1 processing and activation in the vacuole^11, 46, 47^. We found that loss of both *PEP4* and *PPN2* conferred a long-chain phenotype similar to that seen in *ppn1Δ ppn2Δ* cells and countered polyP loss in *sky1Δ* mutants (**Fig. 6A & Fig. 6B**). Interestingly, the *pep4*Δ *ppn2*Δ mutant retained a population of medium length polyP chains absent from the *ppn1*Δ *ppn2*Δ mutant. This effect was exacerbated by alkaline stress and by *SKY1* deletion (**Fig. 6A**) but is not easily explained by changes in total polyP (**Fig. 6B**). We suggest that Ppn1 retains some activity in its unprocessed form or that other proteases can substitute for Pep4 activity under alkaline conditions. Alternatively, it is possible that *PEP4* deletion impacts VTC activity in a manner that favours the synthesis of shorter chains. We propose that increased expression of Pep4 in *sky1*Δ mutants contributes to Ppn1 processing that in turn drives polyP degradation during alkaline stress (see Discussion). Interestingly, like in *sky1*Δ mutants, *npl3*Δ mutants have increased Pep4 levels (**Fig. 6C**), even though these deletions have opposing consequences for polyP accumulation. This discrepancy highlights a complex and multi-factorial response to the regulation of polyP levels during alkaline stress.

## DISCUSSION

Although the enzymes involved in polyP synthesis and degradation have been identified in yeast, the pathways that impact polyP metabolism remain to be fully explored. PolyP levels in *sky1*Δ mutants are remarkably low, suggesting that Sky1 may serve as a particularly important node in polyP regulation.

Based on our results and incorporating findings from the literature, we propose a model (**Fig. 7**) in which the perturbation of the delicate equilibrium between Trk1-mediated potassium influx and Pma1-generated proton gradient in *sky1*Δ promotes membrane depolarization and intracellular alkalinization that is further exacerbated by growth in alkaline media. Previous work has demonstrated that extracellular alkalinization activates the PHO regulon^37, 41^. Working in parallel to mediate the response to high pH stress is a complex network including pathways controlled by cAMP, calcineurin, and the Msn2/4 transcription factors (reviewed by Serra - Cardona *et al.*^37^). Hyperaccumulation of intracellular K+ also depolarizes the plasma membrane^34, 35^, a consequence of which could be the activation of the calcineurin pathway that responds to membrane depolarization events^48, 49^. We speculate that these pathways may be involved in the *sky1*Δ phenotypes presented here. How (and indeed if) PHO induction during alkaline stress contributes to polyP degradation remains unclear. There are over 20 genes that comprise the PHO regulon and these include both positive (*e.g. VTC4*) and negative regulators of polyP accumulation (*e.g. PPN1*, *PPN2*). One possibility is that PHO induction in *sky1*Δ mutants is a response to uniquely low levels of polyP during alkaline stress, rather than a driver of polyP turnover in the vacuole. Oddly, potassium starvation was also shown to induce the expression of several phosphate-related genes, including *PHO5*, *PHO84* and some genes encoding VTC subunits, as well as triggering the depletion of polyP stocks^50, 51^. This suggests that any aberrant fluctuations in intracellular potassium levels have a profound impact on phosphate homeostasis.

**Figure 7.**
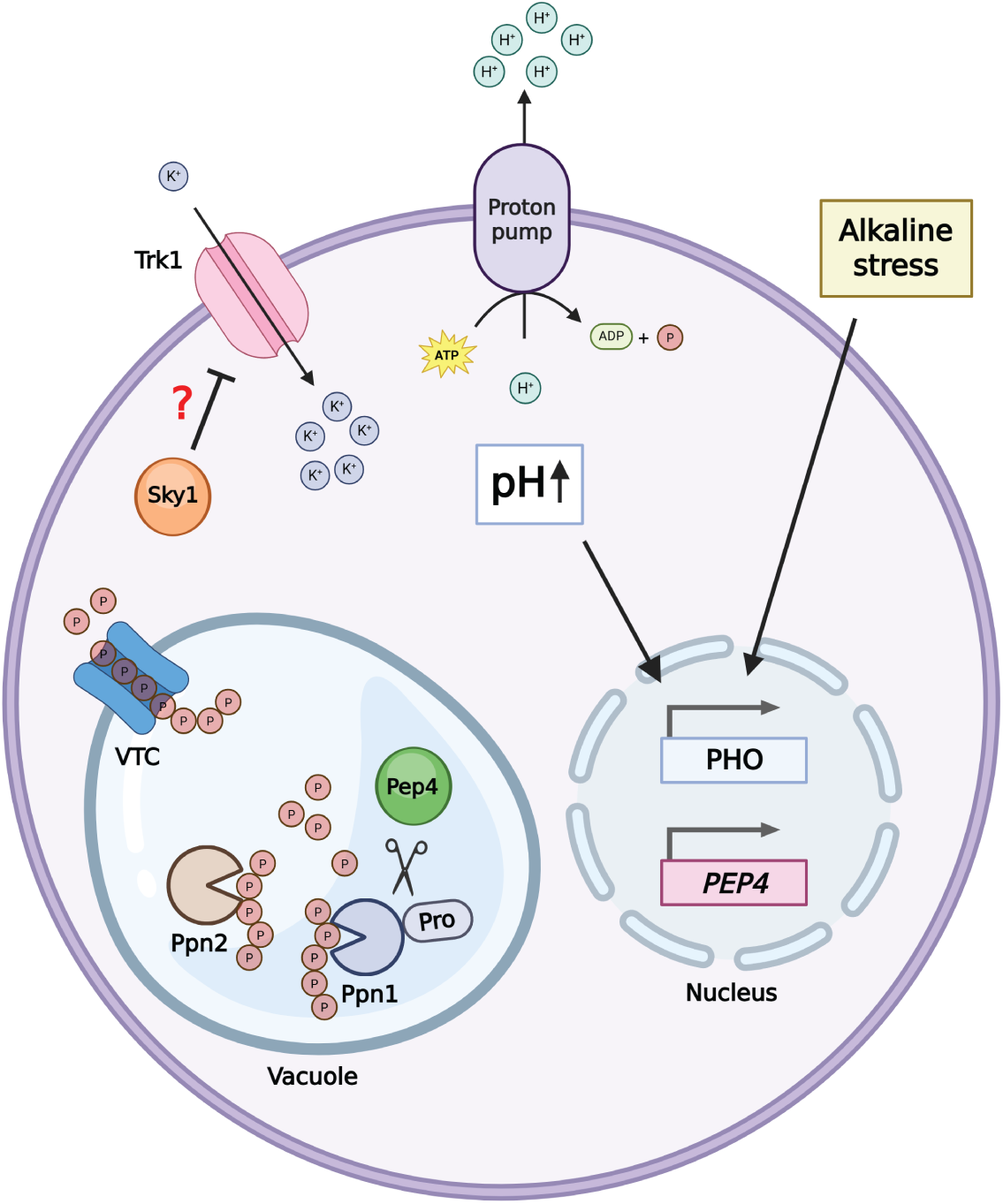
A model for polyP regulation in a *sky1*Δ mutant cell. In the absence of Sky1 potassium ions overaccumulate in the cytosol as the cell fails to properly regulate Trk1 (and Trk2) activity. This prompts increased proton efflux by the Pma1 H+ ATPase to maintain adequate membrane potential, resulting in alkalinization of the cytosol, membrane depolarization and perturbation of the proton electrochemical gradient. That, combined with additional extracellular alkaline stress, triggers the expression of several phosphate-regulated genes via the canonical PHO pathway or in combination with a larger unknown network. Expression of the Pep4 protease is also upregulated in *sky1*Δ mutants, which favors the increased proteolytic activation of its target Ppn1 and, ultimately, the uncontrolled degradation of polyP reserves. Image was made using Biorender.

Regardless of the upstream players, the inability of *sky1*Δ mutants to maintain polyP during alkaline stress appears to be dependent on the action of the Ppn1 and Ppn2 polyphosphatase enzymes residing in the vacuole. As such, we speculate that Ppn1 and/or Ppn2 may be the ultimate effectors of pathways that are regulated by Sky1. Even in the absence of stress, we note that there was no statistically significant difference between the polyP levels in *ppn1*Δ *ppn2*Δ double mutants and *sky1*Δ *ppn1*Δ *ppn2*Δ triple mutants, consistent with a model wherein Sky1 constitutively inhibits the activity of these proteins. A relevant player in this level of control may be the Pep4 protease (**Fig. 7**). Pep4 levels are increased in *sky1*Δ mutants relative to wild-type during alkaline stress and *pep4*Δ *ppn2*Δ mutants largely mimic the polyP phenotypes of *ppn1*Δ *ppn2*Δ counterparts. Vacuolar proteases are required for processing of pro-Ppn1 into an active enzyme and we speculate that in *sky1*Δ mutants increased processing leads to increased Ppn1 action against polyP during alkaline stress. Ppn1 processing is difficult to evaluate in the absence of a high-quality antibody towards the native protein, as the N and C-termini (and any fused epitope tags) are expected to be removed during activation. Production of such a reagent will be critical to properly test our proposed model. Notably, Pep4 has additional vacuolar targets besides Ppn1 that are relevant to phosphate homeostasis, including proteases and the alkaline phosphatase Pho8^52–54^. We speculate that altered processing of these vacuolar proteins could contribute to the *sky1*Δ phenotypes described above. Interestingly, we previously reported that the precursor form of the vacuolar protease Prb1, a Pep4 target, binds to polyP^55^, suggesting the possibility of a complex interplay between polyP maintenance and the regulation of proteolytic activities in the vacuole.

Why Pep4 is misregulated in *sky1*Δ mutants is unclear, but we speculate that *PEP4* transcription in this context could fall under control the same network of cAMP, calcineurin, and Msn2/4 signalling discussed above. Indeed, in wild-type cells Pep4 was previously shown to be strongly and transiently induced 20 minutes after the onset of alkaline stress, and this regulation depended in part on the calcineurin pathway^41^. Beyond polyP turnover, *sky1*Δ cells may experience polyP synthesis defects as VTC activity depends on the proton gradient at the vacuole membrane^3^, which itself relies on the concerted efforts of Pma1 and the vacuolar proton pump V-ATPase^56, 57^.

Our data suggest that Sky1’s kinase activity is required for its role in PHO repression and polyP homeostasis. Whether Sky1 directly phosphorylates Trk1/2 or the phosphatases Ppz1 and Ppz2 is unclear. Rather, Sky1 is thought to preferentially target SR/RS repeats in proteins such as Npl3, Gbp2, and Hrb1. Of these, Npl3 may be most relevant to the phenotypes reported here. Specifically, *sky1*Δ *npl3*Δ double mutants retain at least some polyP after 3 hours at elevated pH, in contrast to *sky1*Δ single mutants. However, our work does not yet support the notion that Sky1 phosphorylation of Npl3 regulates its role in polyP maintenance. The polyP phenotypes of *sky1*Δ and *npl3*Δ mutants are not completely epistatic and expression of Npl3 mutated for a previously characterized Sky1 phosphorylation site did not recapitulate the *sky1*Δ phenotypes. Regardless, we cannot completely rule out a direct role for Sky1 phosphorylation of Npl3 in polyP regulation, since Sky1 may target other sites in addition to S411. It is possible that analysis of additional phosphorylation sites on Npl3 would clarify the relationship between these important regulators and this will be a focus of future work. Further, we remained intrigued by the observation that beyond polyP, *sky1*Δ and *npl3*Δ mutants have opposing effects on the expression of *PHO5* and *PHO84* in response to alkaline stress. Irrespective of its regulation by Sky1, investigation of Npl3 as a novel regulator of phosphate homeostasis and whether it functions upstream, downstream, or in parallel to a Sky1-Trk1 axis is warranted.

Sky1 is a conserved protein kinase, which begs the question of whether its homologs SRPK1-3 impact polyP homeostasis in human cells. While no links to polyP have been reported thus far for SRPKs, we note that members of other SR protein kinase families can interact with polyP. PolyP has been shown to inhibit the activity of CLK3^58^ and DYRK1A^59^ resulting in the modulation of nuclear speckle biogenesis and stability. Lázaro *et al.* also demonstrated via a luciferase-reporter system that ectopic depletion of polyP in HEK293T cells via overexpression of Ppx1 or NUDT3 reduced alternative splicing efficiency^58^. Since Sky1 was first characterized as a regulator of splicing^38^, whether the low polyP levels in *sky1*Δ impact splicing efficiency presents an exciting avenue for future study.

## METHODS

### Yeast strains

The yeast strains used throughout this paper are of the S288C BY4741 background and were generated using standard techniques^60^. These strains are listed in **Supplementary Table S1**. Deletion strains were verified for the correct insertion of the marker and for absence of the target open-reading frame using PCR analyses. Epitope-tagged strains were verified for the correct insertion of the tagging cassette using PCR and fusion-protein expression was confirmed using western blotting.

### Media switch experiments

Overnight cultures were grown at 30 °C in YPD or SC media with pH adjusted to 4.5 with HCl. The next day, cells were diluted to OD_600_ = 0.2 in YPD or SC media at pH 4.5 and grown at 30 °C to OD_600_ = 0.4. Cells were spun down at 3,000 rpm for 5 minutes and pellets were resuspended in YPD or SC media with pH adjusted to 8 with NaOH. 6 (protein preps), 8 (polyP extractions) or 12 (RNA extractions) OD_600_ equivalents of cells were collected at specific time points indicated for each figure.

### Plasmids

Plasmids generated for this work are listed in **Supplementary Table S2** and will be made available from Addgene upon publication. The *SKY1* and *NPL3* genes and their endogenous promoters were generated by Twist Bioscience (San Francisco, USA) and cloned into pAG36. The *NPL3* gene was codon optimized by Twist Bioscience prior to cloning to accommodate for the repetitive nature of the construct. The K187M mutation in pAG36-*SKY1* and S411A mutation in pAG36-*NPL3-*3HA were generated with the QuikChange Lightning Multi Site-Directed Mutagenesis Kit (Agilent). All constructs were sequenced at Plasmidsaurus (USA) or Génome Québec (Montréal, Canada).

### Polyphosphate extractions and gel analyses

The polyphosphate extraction protocol was adapted from Bru *et al.*^61^ and performed as previously described^62^ with modifications. 8 OD_600_ equivalents of cells pelleted in 2-mL screw cap tubes were resuspended in 400 µL of cold LETS buffer (100 mM LiCl, 10 mM EDTA, 10 mM Tris-HCl pH 7.4, 0.2% SDS), 600 µL of equilibrated phenol pH 8 and 200 µL of acid-washed glass beads. Samples were lysed with a BioSpec mini-bead-beater for 2 minutes, heated at 65 °C for 5 minutes and cooled down on ice. 600 µL of chloroform were added followed by 2 minutes of bead beating. Tubes were briefly spun at 1,000 × *g* to clear the lids and a pea-sized amount of high-vacuum silicone grease was added into the lid. Samples were spun down for 2 minutes at 13,000 × *g* and the supernatants were transferred to new 2-mL screw cap tubes containing 600 µL of chloroform and 200 µL of acid-washed glass beads. Samples were lysed as described above and the high-vacuum silicone grease step was repeated a second time. Supernatants were transferred to pre-chilled 1.7-mL microtubes containing 1 mL of 100% ethanol and 40 µL of 3 M sodium acetate pH 5.2. Polyphosphate was precipitated overnight at -20 °C and samples were spun down for 20 minutes at 13,000 × *g* at 4 °C. Supernatants were discarded and 500 µL of 70% ethanol were added, followed by a 5-minute spin at 13,000 × *g* at 4 °C. Supernatants were carefully discarded and precipitated polyP was air dried for 10 minutes before being resuspended in 20 µL of water.

For gel analysis, 5 µL of precipitated polyP was mixed with 9 µL of polyP sample buffer (10 mM Tris-HCl pH 7, 1 mM EDTA, 30% glycerol, traces of bromophenol blue) and loaded onto a 15.8% TBE-urea gel (5.25 g urea, 7.9 mL 30% acrylamide/bis solution (37.5:1), 3 mL 5X TBE buffer, 150 µL 10% ammonium persulfate, 15 µL TEMED). Electrophoresis was performed in 1X TBE buffer at 100 V. Gels were stained in toluidine blue solution (25% methanol, 5% glycerol, 0.05% toluidine blue) for 15 minutes followed by destaining in the same solution without toluidine blue.

### Polyphosphate quantifications

Polyphosphate quantifications were performed using the Phosfinity total polyphosphate quantification kit (Aminoverse) according to the manufacturer’s protocol. Statistical analysis was performed using GraphPad Prism version 10 and significance was determined by ordinary two-way ANOVA with Tukey’s multiple comparisons test.

### Western blotting

Protein extractions were carried out as previously described^62^ with minor modifications, which is reiterated here with similar wording. 6 OD_600_ equivalents of cells were resuspended in 300 µL of 20% trichloroacetic acid (TCA) and 200 µL of acid-washed glass beads. Samples were lysed via bead beating for 2 cycles of 3 minutes and cooled on ice in between cycles. Supernatants were transferred to 1.7-mL tubes and remaining beads were washed in 300 µL of 5% TCA and vortexed. Supernatants were collected and transferred to the previous 1.7-mL tubes. Samples were spun down for 4 minutes at 16,000 × *g* at 4 °C and the supernatants discarded. Pellets were resuspended in 100 µL of 3X sample buffer (160 mM Tris-HCl pH 6.8, 30% glycerol, 6% SDS, and 0.004% bromophenol blue) supplemented with 1/10 volume 1 M dithiothreitol (DTT) and 1/10 volume of 1.5 M Tris-HCl pH 8.8 and heated for 10 minutes at 65 °C. Samples were centrifuged for 4 minutes at 17,000 × *g* and the supernatants were transferred to a new tube. Samples were loaded onto an 8% or 10% SDS-PAGE gel for electrophoresis and separated proteins were transferred to a PVDF membrane for immunoblotting. Membranes were exposed to HyBlot CL autoradiography films (Thomas Scientific). Ponceau S staining or anti-Pgk1 immunoblotting was used as loading control. Antibodies used to detect protein targets, their source, and conditions for use are described in **Supplementary Table S3**.

### RNA extractions

The RNA extraction, cDNA synthesis and RT-qPCR protocol was performed as previously described^62^. 12 OD_600_ equivalents of cells were resuspended in 1 mL of cold TRIzol and 200 µL of acid-washed beads were added. Samples were bead beated for 6 cycles of 30 seconds with 2 minutes on ice in between each cycle. 200 µL of chloroform were added and the samples were vortexed for 20 seconds followed by a 5-minute incubation at room temperature. Cell extracts were centrifuged at 16,000 × *g* for 15 minutes at 4 °C and the top layer was transferred to a new 2-mL screw cap tube containing 400 µL of chloroform followed by 20 seconds of vortexing. Samples were incubated and centrifuged as described above and the top aqueous phase was transferred to a fresh 1.7-mL tube. 400 µL of cold 100% isopropanol and 1 µL of Glycoblue coprecipitant were added and the RNA was precipitated during a 30-minute incubation at -20 °C. Samples were centrifuged at 20,000 × *g* at 4 °C for 20 minutes and the supernatant was discarded. RNA pellets were washed with 1 mL of cold 70% ethanol followed by another 10-minute centrifugation at 20,000 × *g* at 4 °C. Supernatants were discarded and pellets were air dried for 10 minutes in a fume hood. RNA was dissolved in 30 µL of RNase-free water and heated at 55 °C for 5 minutes. After checking the quality of the RNA extractions on an agarose gel, 10 µg of RNA were treated with Ambion DNase I (Invitrogen) according to the manufacturer’s protocol. 100 µL of RNase-free water and 200 µL of phenol-chloroform-isoamyl solution (25:24:1) were added followed by 2 minutes of vortexing. Samples were centrifuged at 20,000 × *g* for 10 minutes at 4 °C and 160 µL of the aqueous phase were transferred to a 1.7-mL tube containing 1 mL of cold 100% ethanol and 16 µL of 3 M sodium acetate pH 5.2. RNA was left to precipitate overnight at -20 °C. Samples were centrifuged for 20 minutes at 20,000 × *g* at 4 °C and supernatants were discarded. Pellets were washed with 1 mL of cold 70% ethanol and RNA was pelleted for 10 minutes at 20,000 × *g* at 4 °C. Supernatants were discarded and pellets were air dried for 10 minutes in a fume hood. RNA was dissolved in 25 µL of RNase-free water followed by a 5-minute incubation at 55 °C.

### cDNA synthesis and RT-qPCR analyses

1 µg of RNA was transcribed to cDNA using 5× All-In-One reverse transcription (RT) master mix (Applied Biological materials). The thermocycler conditions were 37 °C for 15 minutes, 25 °C for 10 minutes, 60 °C for 50 minutes and 85 °C for 5 minutes. The cDNA was diluted 1:5 in RNase-free water for subsequent analyses. qPCRs were performed using iQ SYBR green supermix (Bio-Rad) according to the manufacturer’s protocol in 3 technical replicates with ≥ 3 biological replicates. The thermocycler conditions were 95 °C for 3 minutes and 40 cycles of 95 °C for 15 seconds, 60 °C for 30 seconds, and 72 °C for 30 seconds with melt curve analyses performed after each run. Standard curves using genomic DNA were included to assess efficiency of each primer pair which are listed in **Supplementary Table S4**. Changes in mRNA expression were normalized with *TAF10* as housekeeping gene and calculated using the ΔΔ*C_T_* method with the Bio-Rad CFX Maestro 2.3 version 5.3.022.1030 software. Statistical analysis was performed using GraphPad Prism version 10 and significance was determined by ordinary one-way ANOVA with Tukey’s multiple comparisons test.

### Statistical analyses

Statistical analyses are described separately in the relevant sections of the methods.

## ACKNOWLEDGEMENTS

We acknowledge members of the Downey lab for critical reading of the manuscript and for valuable suggestions. This work was funded by a Canadian Institutes of Health Research (CIHR) grant to MD (PJT-525398). SM was supported by a Natural Sciences and Engineering Research Council of Canada (NSERC) USRA award. We thank Scott Emr for providing the Pep4 antibody used in this work.

---

**Figure S1.**
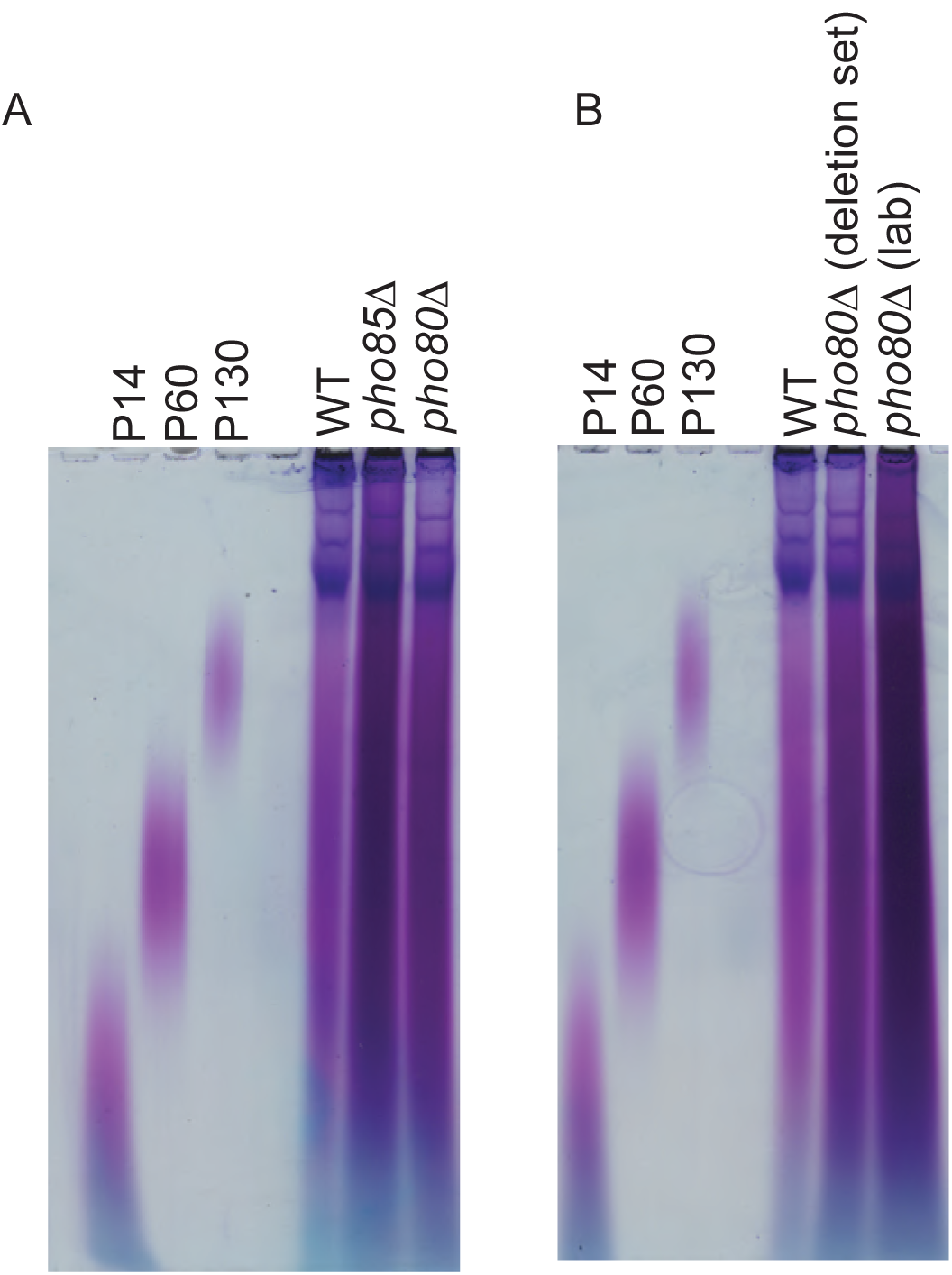
PolyP phenotype of lab-made *pho80*Δ strain differs from that of the yeast deletion set. **(A)** and **(B)** PolyP was extracted from cells grown to exponential phase in YPD media and stained with toluidine blue following TBE-urea PAGE separation. Image is representative of data from ≥ 3 experiments.

**Figure S2.**
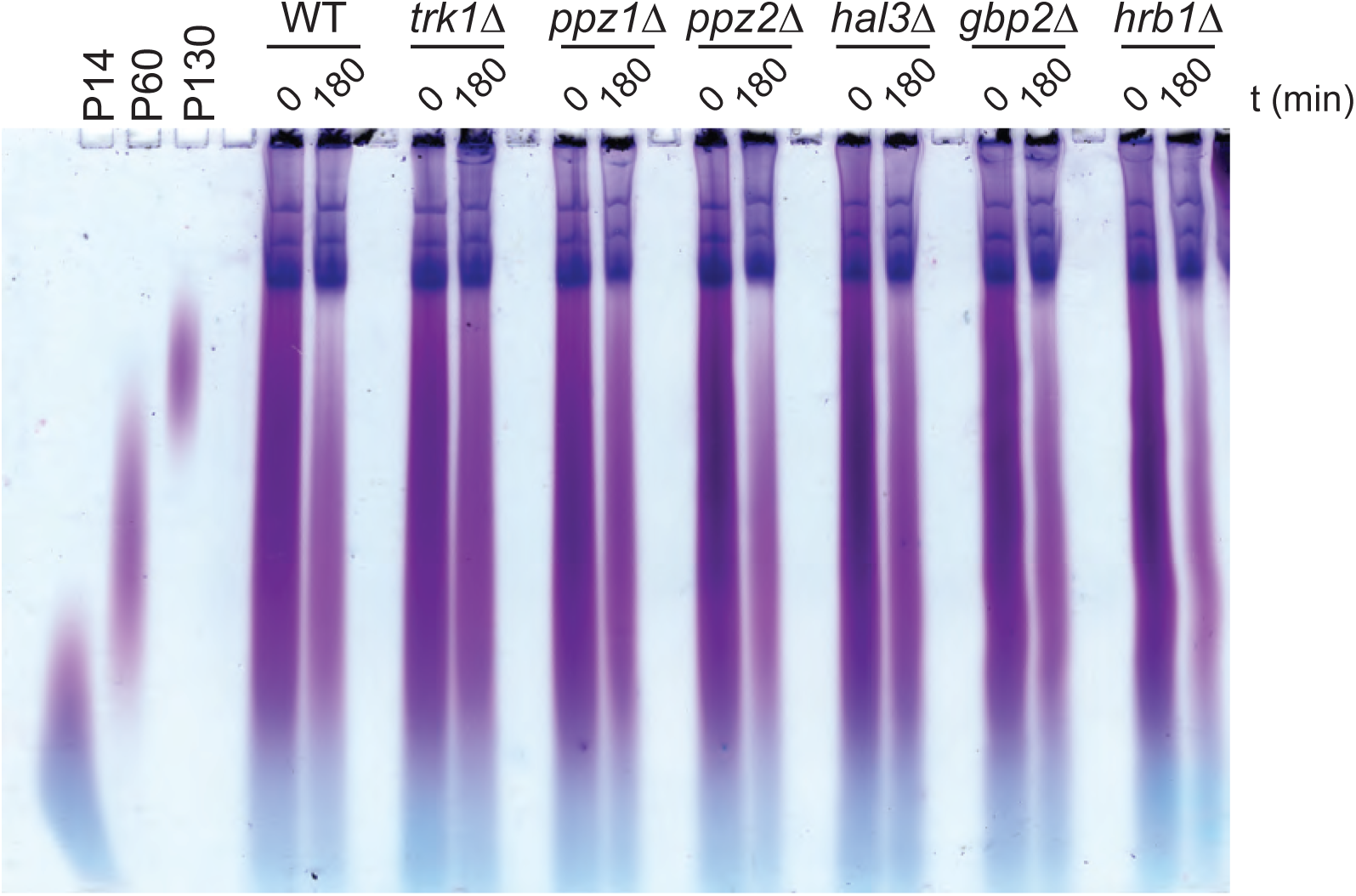
PolyP analyses for single mutants used in this study. Indicated strains were grown in YPD pH 4.5 for approximately 1 cell division then switched to YPD pH 8. Cells were collected before the switch and after 3 hours in alkaline conditions, and polyP was extracted for TBE-urea PAGE separation and toluidine blue staining. Image is representative of data from ≥ 3 experiments.

**Figure S3.**
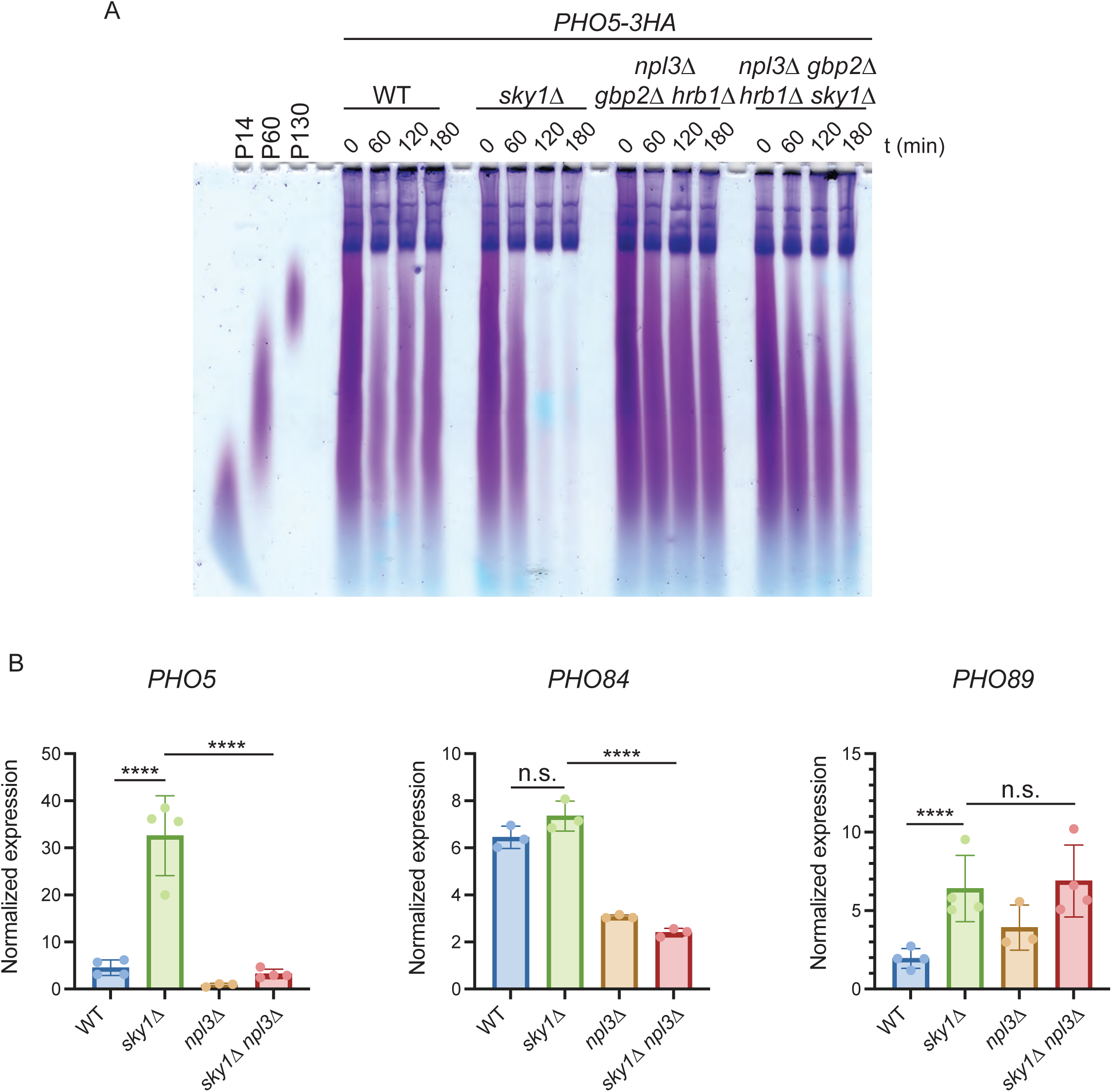
Triple deletion of *NPL3*, *GBP2* and *HRB1* increases polyP levels similarly to *npl3*Δ mutants. **(A)** Indicated strains were grown in YPD pH 4.5 for approximately 1 cell division then switched to YPD pH 8. Cells were collected at 1 hour intervals and polyP was extracted for TBE-urea PAGE separation and toluidine blue staining. Image is representative of data from ≥ 3 experiments. **(B)** *PHO5*, *PHO84* and *PHO89* mRNA analysis. Strains were grown in YPD pH 4.5 for approximately 1 cell division then switched to YPD pH 8. Cells were collected after 3 hours in alkaline conditions and RNA was extracted for RT-qPCR analyses performed in triplicates. Data are presented as means ± SD. Significance was determined by ordinary one-way ANOVA based on Log2 of normalized expression data with Tukey’s multiple comparisons test. ****, *p* < 0.0001. n.s., non-significant. Significance is indicated only for key comparisons discussed in the text, with all *p* values presented in **Supplementary Table S5**.

**Figure S4.**
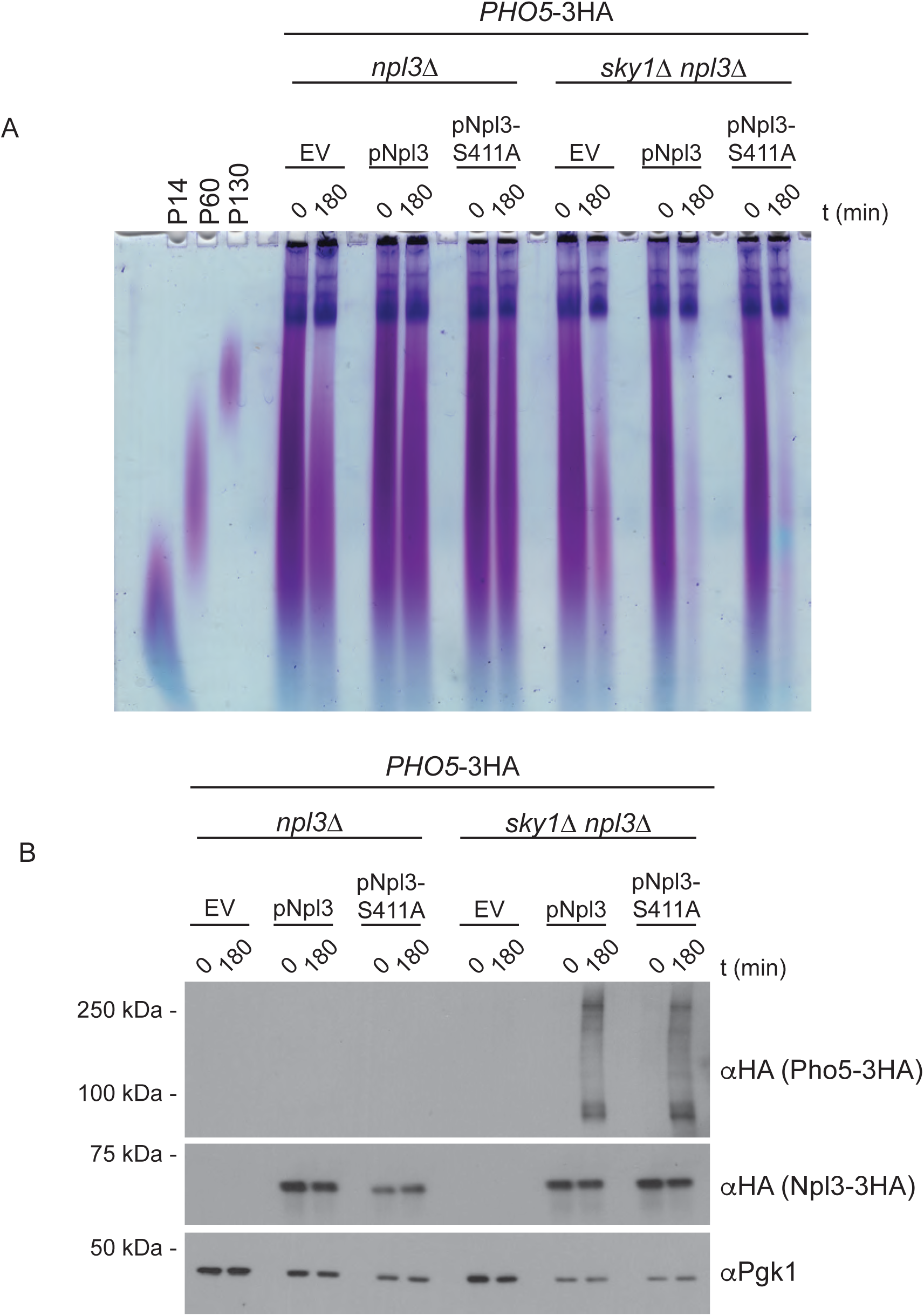
Mutation of S411 to alanine on Npl3 does not replicate *sky1*Δ’s polyP and Pho5-3HA phenotypes. **(A)** and **(B)** *PHO5*-3HA strains were transformed with an empty vector or a vector expressing codon optimized *NPL3*-3HA or *NPL3* S411A-3HA under its native promoter. Cells were grown in YPD pH 4.5 for approximately 1 cell division then switched to YPD pH 8. Cells were collected before the switch and after 3 hours in alkaline conditions for polyP **(A)** or protein **(B)** extractions and PAGE analysis. PolyP gels were stained with toluidine blue. Proteins transferred onto PVDF membrane were probed with an anti-HA antibody, and an anti-Pgk1 antibody was used as loading control. Images are representative of data from ≥ 3 experiments.

**Figure S5.**
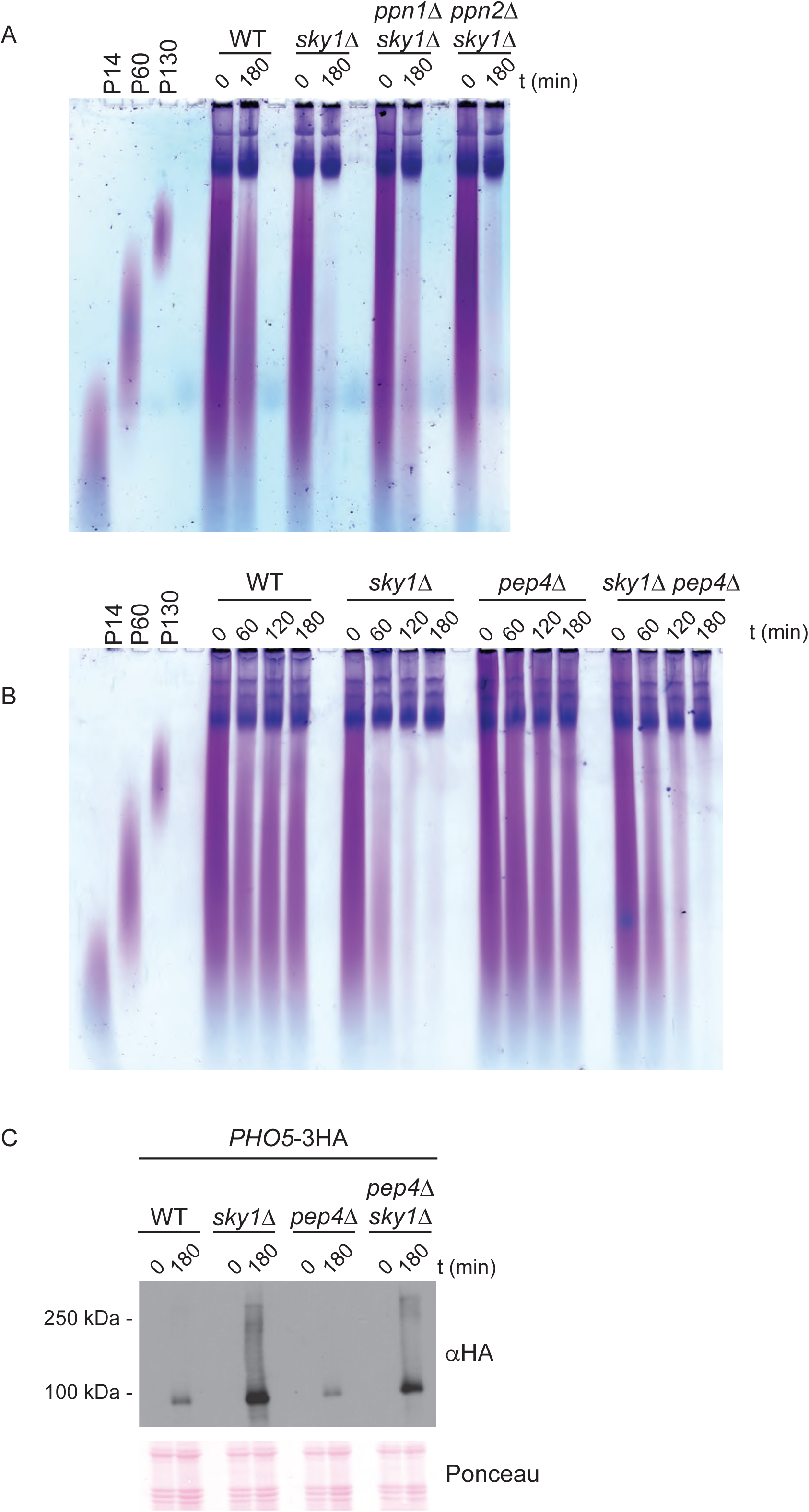
Single deletion of *PPN1* or *PPN2* in *sky1Δ* mutants does not prevent the loss of polyP in alkaline conditions. **(A)** Indicated strains were grown in YPD pH 4.5 for approximately 1 cell division then switched to YPD pH 8. Cells were collected before the switch and after 3 hours in alkaline conditions, and polyP was extracted for TBE-urea PAGE separation and toluidine blue staining. **(B)** Indicated strains were grown as above with additional time points after the switch to YPD pH 8. PolyP was extracted and analyzed as above. **(C)** Indicated strains were grown as in **(A)** and cells were collected for protein extraction, SDS-PAGE separation and western blotting. Membrane was probed with an anti-HA antibody and Ponceau S staining of the membrane is shown as loading control. Images are representative of data from ≥ 3 experiments.

**Supplementary Table S1. – Yeast strains used in this study**

**Supplementary Table S2. – Plasmids used in this study**

**Supplementary Table S3. – Antibodies used in this study**

**Supplementary Table S4. – Oligonucleotides used in this study**

**Supplementary Table S5. – *p* values for all statistical tests**

